# PE5-PPE4-EspG_3_ trimer structure from mycobacterial ESX-3 secretion system gives insight into cognate substrate recognition by ESX systems

**DOI:** 10.1101/2020.01.16.909523

**Authors:** Zachary A. Williamson, Catherine T. Chaton, William A. Ciocca, Natalia Korotkova, Konstantin V. Korotkov

**Author notes:** To whom correspondence should be addressed: Konstantin V. Korotkov: Department of Molecular & Cellular Biochemistry, University of Kentucky, Lexington, KY 40536. Department of Biological Sciences, Eastern Kentucky University, Richmond, KY 40475.

## Abstract

*Mycobacterium tuberculosis* (*Mtb*) has evolved numerous type VII secretion (ESX) systems to secrete multiple factors important for both growth and virulence across their cell envelope. Three such systems; ESX-1, ESX-3, and ESX-5; have been shown to each secrete a unique set of substrates. A large class of these substrates secreted by these three systems are the PE and PPE families of proteins. Proper secretion of the PE-PPE proteins requires the presence of EspG, with each system encoding its own unique copy. There is no cross-talk between any of the ESX systems and how each EspG is recognizing its subset of PE-PPE proteins is currently unknown. The only current structural characterization of PE-PPE-EspG trimers is from the ESX-5 system. Here we present the crystal structure of the PE5_mt_-PPE4_mt_-EspG_3mm_ trimer, from the ESX-3 system. Our trimer reveals that EspG_3mm_ interacts exclusively with PPE4_mt_ in a similar manner to EspG_5_, shielding the hydrophobic tip of PPE4_mt_ from solvent. The C-terminal helical domain of EspG_3mm_ is dynamic, alternating between an ‘open’ and ‘closed’ form, and this movement is likely functionally relevant in the unloading of PE-PPE heterodimers at the secretion machinery. In contrast to the previously solved ESX-5 trimers, the PE-PPE heterodimer of our ESX-3 trimer is interacting with it’s chaperone at a drastically different angle, and presents different faces of the PPE protein to the chaperone. We conclude that the PPE-EspG interface from each ESX system has a unique shape complementarity that allows each EspG to discriminate amongst non-cognate PE-PPE pairs.

---

Tuberculosis is currently the deadliest infectious disease in the world, killing 1.5 million people in 2019 (1). The lack of an effective vaccine against the most prevalent pulmonary form of Tuberculosis, as well as the emergence of numerous multi-drug resistant strains of the causative agent, *Mycobacterium tuberculosis* (*Mtb*), highlights the growing need for more effective treatment options. Therefore, a more comprehensive understanding of the *Mtb* virulence machinery is needed to aid the development of new therapeutics.

*Mtb*, like all mycobacteria, contains a thick hydrophobic cell envelope that aids in protecting the mycobacterium from its environment. To overcome the limited permeability created by this envelope, mycobacteria have evolved a specialized secretion system to export proteins across their cell envelopes, the type VII secretion system, also known as the ESX system (2). Five different ESX systems encoded in the *Mtb* genome, and three are known to secrete proteins; ESX-1, ESX-3, and ESX-5 (3). Recently, the structures of the core complex of both ESX-3 (4,5) and ESX-5 (6) have been solved. The ESX-5 core complex has six-fold symmetry and sits on the inner membrane (6), while the ESX-3 core complex was solved as a dimer that could be modeled onto the six-fold symmetry of the ESX-5 core complex (4,5). These systems are not functionally redundant, as their substrates are not re-routed to other ESX systems (7). The ESX systems secrete a variety of different substrates, each containing a general type VII secretion motif of YxxxD/E (8). A significant class of substrates being the PE and PPE proteins, named for conserved residues (Pro-Glu for PE and Pro-Pro-Glu for PPE) within their N-terminal domains (9,10). The N-terminal domains are about 110 (PE) or 180 (PPE) amino acids in length and interact together to form a PE-PPE heterodimer. A cytosolic chaperone, EspG, is required for proper folding and/or stability of the PE-PPE proteins and, ultimately their proper secretion (11,12). Each ESX system secretes a unique subset of PE-PPE heterodimers, and therefore each encodes an EspG that binds to only its corresponding heterodimers (11,12). The first structural insight into the EspG and PE-PPE interaction was revealed by analysis of the structure of the PE25-PPE41-EspG_5_ complex, a trimer from ESX-5 (12,13). EspG_5_ interacts solely with PPE41 at the tip distal to the PE25 interaction and aids in preventing PE-PPE heterodimer aggregation in part by shielding a conserved hydrophobic tip on the PPE proteins, known as the hh motif (12). The additional structure of the ESX-5-related, PE8-PPE15-EspG_5_ trimer, revealed similar interactions of the substrate PE-PPE dimer with the EspG_5_ chaperone (14). Despite high conservation among PPE proteins in the identified EspG_5_ binding region from PPE41, three residues vary depending on whether the PPE protein is secreted by ESX-1, ESX-3, or ESX-5 (12). Alteration of any or all of these positions in the ESX-5-dependent PPE41 did not disrupt PPE41-EspG_5_ binding (12). Based on this observation it has been suggested that structural elements outside of the EspG-binding region differentiate the ESX-5-specific PPE proteins from their ESX-1 and ESX-3 homologs to bind EspG_5_ (12).

This study was initiated to understand the how each EspG from the different ESX systems specifically recognizes its unique subset of cognate PE-PPE heterodimers. Here we present the structure of PE5-PPE4-EspG_3_ from ESX-3. This structure reveals a novel binding mode of PE-PPE proteins with the EspG chaperone and suggests the molecular mechanism by which the PE-PPE dimers are specifically targeted by cognate chaperones.

## Results

### EspG_3_ forms a complex with PE5-PPE4 and binding is conserved across species

In order to understand the mechanism for the specificity of PE-PPE recognition by cognate chaperones a high-resolution structure of a trimer produced by the ESX systems, other than ESX-5, was needed. Our efforts have focused on optimizing the ESX-3 PE-PPE-EspG trimer for X-ray structural studies. Constructs of full-length PE5 (*Rv0285*), the conserved N-terminal PPE domain of PPE4 (*Rv0286*, residues 1-181), in a complex with the cognate full-length EspG_3_ (*Rv0289*) from *Mycobacterium tuberculosis* (Fig. 1*a*) never formed high-resolution diffraction quality crystals, despite our best efforts. The difficulty could be due to some heterogeneity in the processing of EspG_3mt_ within the *Escherichia coli* cell, as seen by the double band in Fig. 1*b* and Sup. Fig. 1*a*. Numerous variations of PE5-PPE4-EspG_3_ constructs were screened utilizing multiple mycobacterial species, different fusion approaches, and even mixing PE5-PPE4 dimers with EspG_3_ chaperones from different species (Sup. Table 1). This latter approach was inspired by the work done on the *Plasmodium* aldolase-thrombospondin-related anonymous protein complex (15), and in the end, produced the best crystals for further diffraction experiments. To ensure the mixed trimers behaved the same in solution as the wild-type (WT) trimer, size-exclusion chromatography with multi-angle light scattering (SEC-MALS) experiment was performed on both the WT PE5_mt_-PPE4_mt_-EspG_3mt_ trimer as well as the mixed PE5_mt_-PPE4_mt_-EspG_3mm_ trimer which contained the *Mycobacterium marinum* EspG_3_ gene (*MMAR_0548*) (Fig. 1*b-c*). Both trimers form a 1:1:1 complex with experimental molecular weights of 56.2 kDa (Fig. 1*b*) for the full *M. tuberculosis* trimer and 54.6 kDa (Fig. 1*c*) for the mixed trimer with the *M. marinum* EspG_3_. Co-purification assays were run with both *M. tuberculosis* and *M. marinum* EspG_3_ with the *M. tuberculosis* PE4-PPE5, along with EspG_3_’s from *Mycolicibacterium smegmatis (MSMEG_0622), Mycolicibacterium hassiacum* (*MHAS_04631*), and *Mycobacterium kansasii* (*MKAN_17015*). Due to the His_6_-tag only being present on PE5_mt_, EspG_3_ co-purification required interaction with the PE5_mt_-PPE4_mt_ heterodimer. Across all species that were tested, EspG_3_ co-purified with the PE5_mt_-PPE4_mt_ heterodimer (Supplementary Fig. 1*a-e*). The binding of different EspG_3_s to the same PE-PPE heterodimer suggests a common protein-protein recognition mechanism within the ESX-3 family.

**Figure 1.**
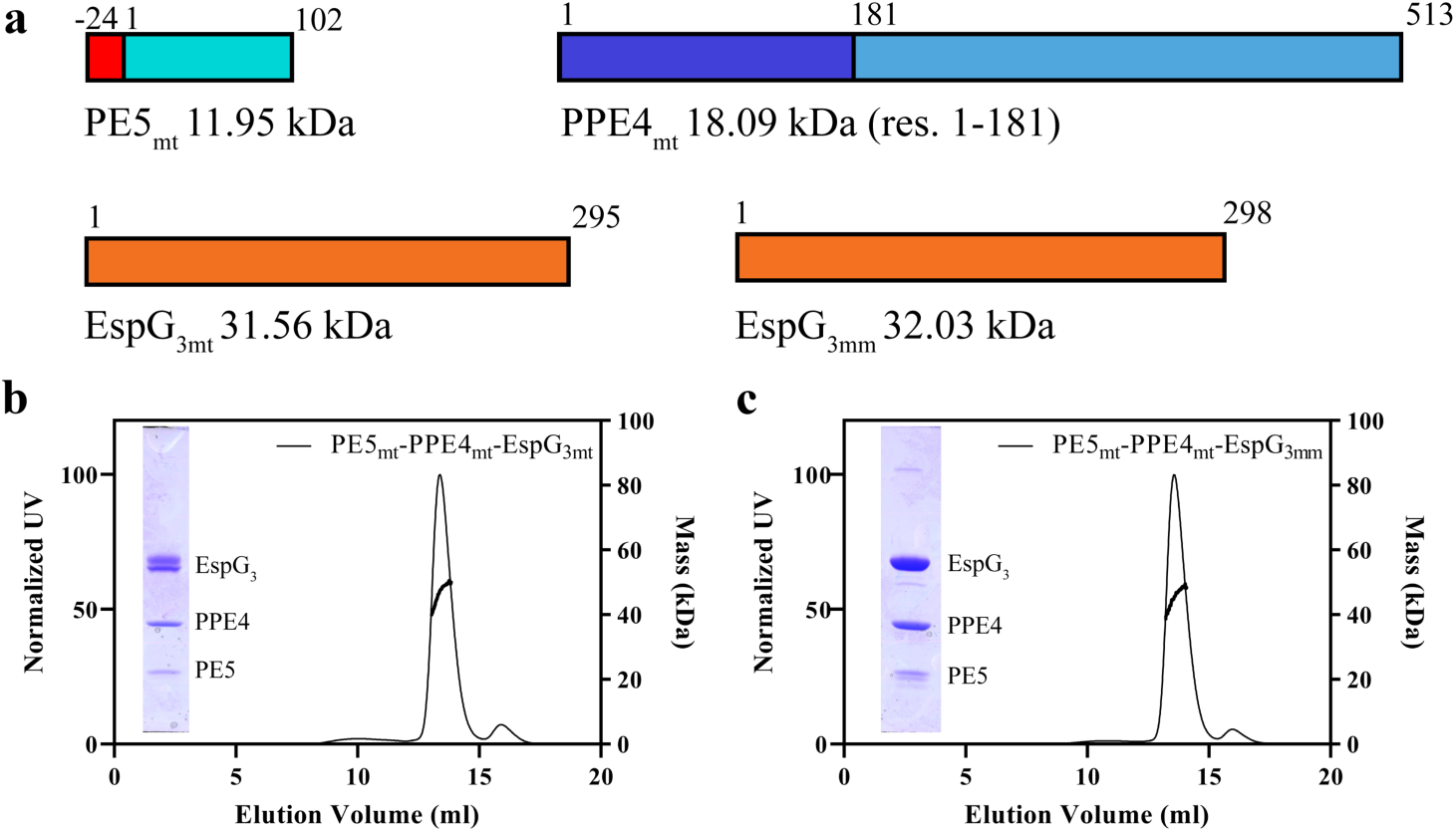
Solution characterization of the PE5-PPE4-EspG_3_ trimer. *a*, schematic showing design and molecular weights for constructs used in this study. PE5 from *M. tuberculosis* (*Rv0285*) contains an N-terminal His_6_-tag that is connected to the gene via a TEV protease cleavable linker. PPE4 (*Rv0286*) was truncated after its N-terminal PPE domain. Full-length copies of both *M. tuberculosis* (*Rv0289*) and *M. marinum* (*MMAR_0548*) EspG_3_ were used. *b* and *c*, elution profile of PE5_mt_-PPE4_mt_-EspG_3mt_ (*b*) and PE5_mt_-PPE4_mt_-EspG_3mm_ (*c*), with the right Y-axis showing the MALS-measured molecular weight. Inset shows an SDS-PAGE image of the peak fraction.

### Overall structure of PE5_mt_-PPE4_mt_-EspG_3mm_

The PE5_mt_-PPE4_mt_-EspG_3mm_ trimer was able to form diffraction quality crystals, and two different crystal forms were observed that diffracted to 3.3 Å (I422) and 3.0 Å (P2_1_2_1_2_1_) (Table 1). Final refinement and data statistics are in Table 1. Overall there is little structural variation between the individual proteins across the copies present in the two crystal forms (Table 2).

**Table 1.** Data collection and refinement statistics.

|  | PE5<sub>mt</sub>-PPE4<sub>mt</sub>-EspG<sub>3mm</sub><br>(6UUJ) | PE5<sub>mt</sub>-PPE4<sub>mt</sub>-EspG<sub>3mm</sub><br>(6VHR) |
| --- | --- | --- |
| **Data Collection** |  |  |
| Wavelength (Å) | 1.000 | 1.000 |
| Space group | P2<sub>1</sub>2<sub>1</sub>2<sub>1</sub> | I422 |
| Cell Dimensions: |  |  |
| *a*, *b*, *c* (Å) | 72.26, 158.63, 209.31 | 219.14, 219.14, 104.44 |
| $\alpha$, $\beta$, $\gamma$(°) | 90, 90, 90 | 90, 90, 90 |
| Resolution (Å) | 39.51 – 3.00 (3.08 – 3.00)<sup>a</sup> | 35.73 – 3.30 (3.39 – 3.30) |
| R<sub>sym</sub> | 0.131 (1.56) | 0.087 (2.18) |
| CC<sub>1/2</sub><sup>b</sup> | 0.998 (0.590) | 0.999 (0.597) |
| *I*/$\sigma$ | 9.45 (1.14) | 15.70 (1.31) |
| Completeness (%) | 99.1 (99.5) | 99.8 (100) |
| Multiplicity | 4.2 (4.4) | 10.3 (9.5) |
| **Refinement** |  |  |
| Resolution (Å) | 39.51 – 3.00 | 35.73 – 3.30 |
| No. reflections (total/free) | 48463/2462 | 19335/928 |
| R<sub>work</sub>/R<sub>free</sub> | 0.266/0.303 | 0.248/0.266 |
| Number of atoms: |  |  |
| Protein | 14535 | 3643 |
| Ligand/ion | 0 | 0 |
| Water | 4 | 0 |
| *B*-factors: |  |  |
| Protein | 101.6 | 173.2 |
| Water | 70.6 |  |
| All atoms | 101.6 | 173.2 |
| Wilson *B* | 87.9 | 147.3 |
| R.m.s. deviations: |  |  |
| Bond lengths (Å) | 0.002 | 0.002 |
| Bond angles (°) | 0.53 | 0.503 |
| Ramachandran distribution<sup>c</sup> (%) |  |  |
| Favored | 96.42 | 91.34 |
| Allowed | 3.58 | 8.04 |
| Outliers | 0 | 0.62 |
| Rotamer outliers<sup>c</sup> (%) | 0.28 | 0 |
| Clashscore<sup>d</sup> | 7.02 | 6.94 |
| MolProbity Score<sup>e</sup> | 1.62 | 1.89 |

<sup>a</sup>Values in parentheses are for the highest resolution shell.<sup>b</sup>CC<sub>1/2</sub> correlation coefficient is defined in (Karplus and Diederichs) and was calculated with *XSCALE* (Kabsch).<sup>c</sup>Calculated with the MolProbity server (http://molprobity.biochem.duke.edu) (Chen or Williams).<sup>d</sup>Clashscore is the number of serious steric overlaps (> 0.4 Å) per 1000 atoms.<sup>e</sup>MolProbity Score combines the clashscore, rotamer, and Ramachandran evaluations into a single score, normalized to be on the same scale as X-ray resolution (Chen or Williams)

**Table 2.** Structural Variations in copies of PE5_mt_-PPE4_mt_-EspG_3mm_ structure in r.m.s.d. (Å).

|  | PE5<sub>mt</sub> | PPE4<sub>mt</sub> | EspG<sub>3mm</sub> |
| --- | --- | --- | --- |
| Aligned to 6UUJ copy 1 |  |  |  |
| 6UUJ copy 2 | 0.2 | 0.3 | 0.4 |
| 6UUJ copy 3 | 0.4 | 0.2 | 0.4 |
| 6UUJ copy 4 | 0.3 | 0.2 | 0.4 |
| 6VHR | 0.4 | 0.5 | 0.6 |
| Aligned to 6UUJ copy 2 |  |  |  |
| 6UUJ copy 1 | 0.2 | 0.3 | 0.4 |
| 6UUJ copy 3 | 0.3 | 0.3 | 0.3 |
| 6UUJ copy 4 | 0.3 | 0.3 | 0.4 |
| 6VHR | 0.4 | 0.5 | 0.6 |
| Aligned to 6UUJ copy 3 |  |  |  |
| 6UUJ copy 1 | 0.4 | 0.2 | 0.4 |
| 6UUJ copy 2 | 0.3 | 0.3 | 0.3 |
| 6UUJ copy 4 | 0.4 | 0.3 | 0.4 |
| 6VHR | 0.5 | 0.5 | 0.6 |
| Aligned to 6UUJ copy 4 |  |  |  |
| 6UUJ copy 1 | 0.3 | 0.2 | 0.4 |
| 6UUJ copy 2 | 0.3 | 0.3 | 0.4 |
| 6UUJ copy 3 | 0.3 | 0.3 | 0.4 |
| 6VHR | 0.4 | 0.5 | 0.7 |

For all structural analysis and comparisons, the first copy of the PE5_mt_-PPE4_mt_-EspG_3mm_ trimer from the higher resolution P2_1_2_1_2_1_ crystal form was used because it diffracted at a higher resolution and has the lowest *B*-factors from the non-crystallographic copies in the P2_1_2_1_2_1_ form. EspG_3mm_ interacts solely with the tip of PPE4_mt_ (Fig. 2), similar to EspG_5_ in the previously solved ESX-5 trimers (12-14). However, the orientation of PE5_mt_-PPE4_mt_ relative to EspG_3mm_ is dramatically different than was observed for either ESX-5 trimer, and the differences between them will be described in later sections. The YXXD/E motif for ESX secretion of PE5_mt_ is accessible for interactions with the rest of the ESX machinery, as it is located distal to the EspG_3mm_ interaction (11). In both crystal forms, this secretion motif is disordered, similar to the motif in PE8_mt_ from the PE8_mt_-PPE15_mt_-EspG_5mt_ trimer (14). The individual components of the PE5_mt_-PPE4_mt_-EspG_3mm_ trimer align well to the individual components of the previously reported ESX-5 trimers, both PE25_mt_-PPE41_mt_-EspG_5mt_ (4KXR and 4W4L) and PE8_mt_-PPE15_mt_-EspG_5mt_ (5XFS), with only moderate variations (Table 3).

**Table 3.** Structural variations between individual components of ESX-3 trimer and the previously published ESX-5 trimers in r.m.s.d. (Å)

|  | PE25<sub>mt</sub>-PPE41<sub>mt</sub>-EspG<sub>5mt</sub><br>(4KXR) | PE25<sub>mt</sub>-PPE41<sub>mt</sub>-EspG<sub>5mt</sub><br>(4W4L) | PE8<sub>mt</sub>-PPE15<sub>mt</sub>-EspG<sub>5mt</sub><br>(5XFS) |
| --- | --- | --- | --- |
| PE5<sub>mt</sub> | 2.3 | 2.3 | 1.4 |
| PPE4<sub>mt</sub> | 3.3 | 3.4 | 2.5 |
| EspG<sub>3mm</sub> | 2.4 | 2.7 | 2.3 |

**Figure 2.**
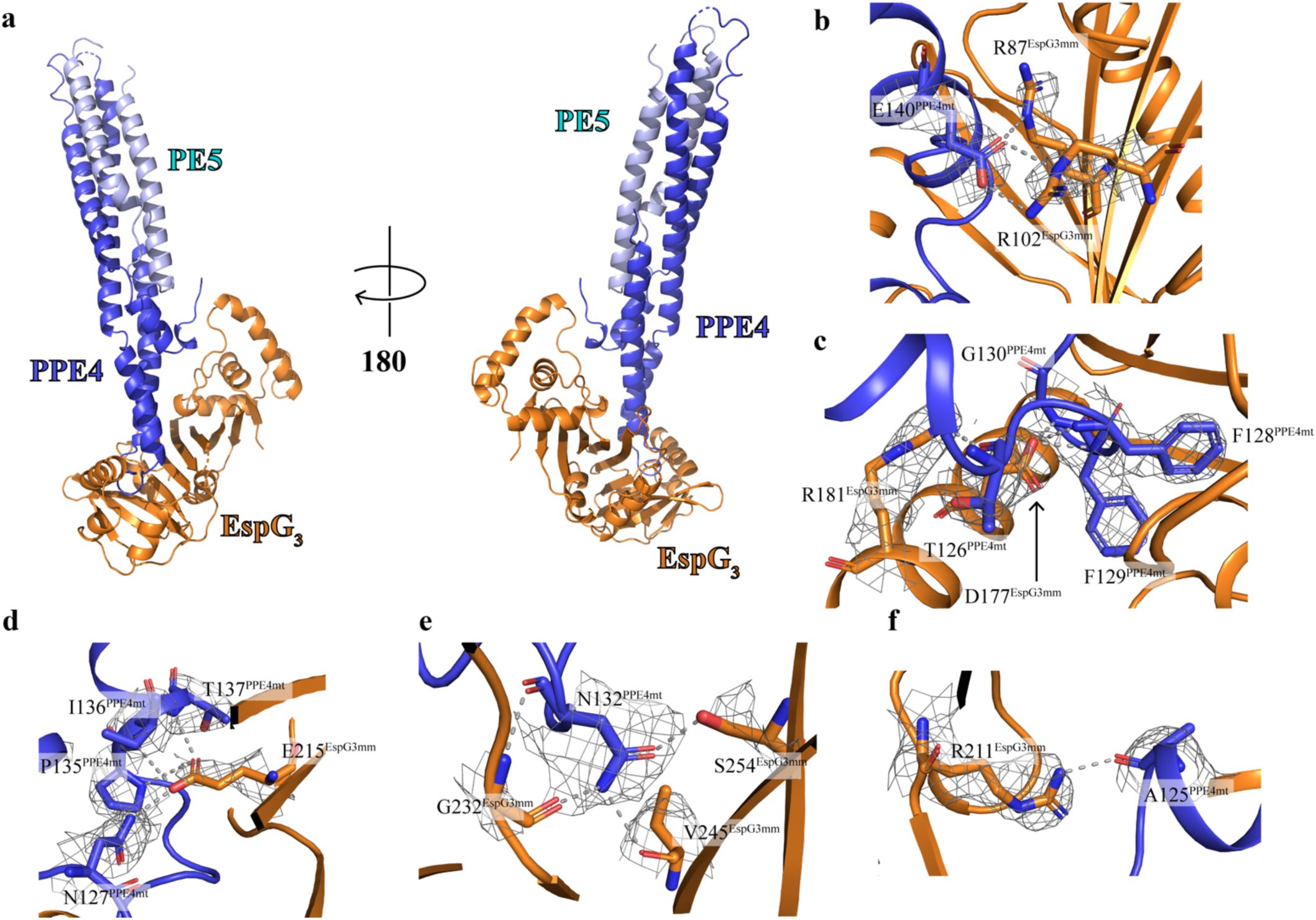
Crystal structure of the PE5_mt_-PPE4_mt_-EspG_3mm_ trimer and selected interactions in PPE4_mt_-EspG_3mm_ interface. *a*, Crystal structure of the PE5_mt_-PPE4_mt_-EspG_3mm_ trimer in a cartoon representation with two views related by a rotation of 180°. EspG_3_ interacts exclusively with the tip PPE4, distal to PE5. *b-f*, Interacting residues are shown with main chain and side chain in stick form with electron density map (2Fo-Fc shown at 1.0 σ) covering side chains and hydrogen bonding in grey dashed lines.

In a previous study on EspG structures (16), a small-angle X-ray scattering (SAXS) experiment was done on the PE5-PPE4-EspG_3_ trimer from *M. smegmatis*. Comparisons between this SAXS analysis and our crystal structure were performed to see if the solution-based characterization of the trimer matched the X-ray-based characterization. We ran CRYSOL (17) on our crystal structure in comparison to the experimental scattering data from the *M. smegmatis* trimer. The overall χ^2^ is 2.53, which is acceptable given that the trimers are from different species with only 54.0-73.8% sequence identity across the different components (Sup. Fig. 2). The main differences are in the extreme high- and low-resolution areas, likely arising from differences in the primary structure between the two samples and from aggregation in the SAXS sample, respectively. Therefore, we are confident that the crystal structure is an appropriate model of the ESX-3 trimer as it exists in solution.

### Interface between PPE4_mt_ and EspG_3mm_

The interface between EspG_3mm_ and PPE4_mt_ contains numerous hydrophobic interactions, multiple hydrogen bonds, and two salt bridges centered around E140^PPE4mt^ (Fig. 2*b-f*). Overall the interface buries 3,121 Å^2^ of solvent-accessible surface area, as calculated by the PISA server (18) and is comprised of 30 total residues from PPE4_mt_ and 49 residues from EspG_3mm_ (Fig. 3). The tip of PPE4_mt_ containing the ends of α4 and α5 and the loop between them is inserted into a groove on EspG_3_ composed of its central β sheet and C-terminal helical bundle. This bundle shields the hydrophobic tip of PPE4_mt_, including the hh motif of FF128, from solvent access. The tip of PPE4_mt_ is interacting with EspG_3_ in such a way that the complex is unlikely to disengage at the ESX secretion machinery without structural rearrangement of the chaperone.

**Figure 3.**
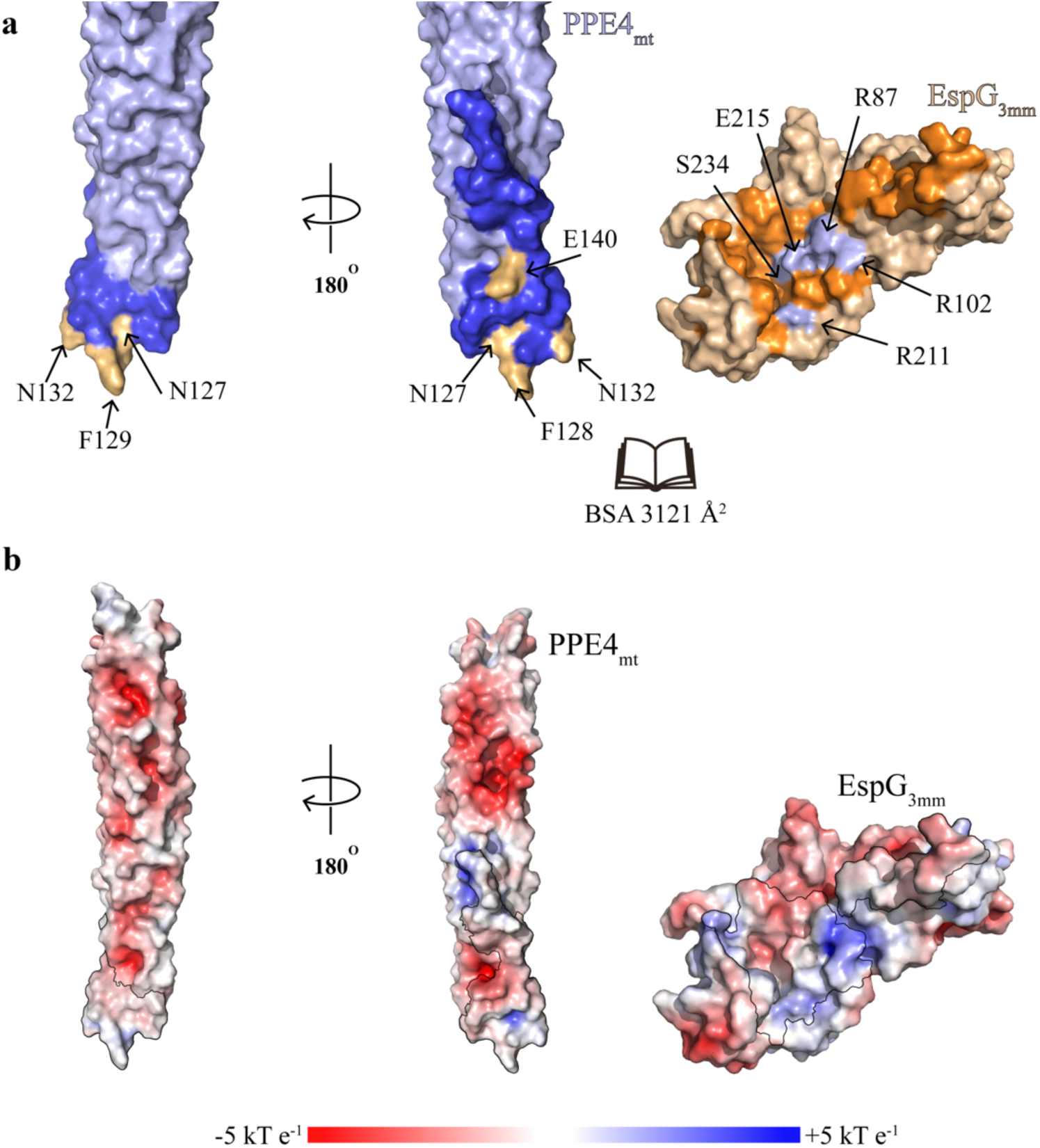
Interface between PPE4_mt_ and EspG_3mm_. *a*, Surface representation of PPE4_mt_ and EspG_3mm_ shown in an ‘open book’ view. Interacting residues are colored in blue (PPE4_mt_) and orange (EspG_3mm_), with mutated residues highlighted in light orange (PPE4_mt_) and light blue (EspG_3mm_). Another view of PPE4_mt_, related by a 180° rotation, is also shown. *b*, The same orientations as panel *a* but with the surface colored according to surface potential as calculated by APBS (ref). The interacting residues are circled within a black line.

### Mutations cause disruptions in the PPE4-EspG_3_ interface

In order to probe the interface of the crystal structure and test the importance of interacting residues, we made several mutations on both PPE4 and EspG_3_ sides of the interface and opted to use the cognate PE5_mt_-PPE4_mt_-EspG_3mt_ trimer to test our mutations. The PISA output (18) of the interface was analyzed along with sequence alignments of the current known ESX-3 PPE proteins (Sup. Fig. 3) and alignments of the EspG_3_ used in this study (Sup. Fig. 4) to select which residues in the interface would be mutated.

PPE4_mt_ is well conserved along the interface among ESX-3-specific PPE proteins (Sup. Fig. 3), and we targeted strictly conserved residues in the interface. We selected N127 and N132 because they contain buried hydrogen bonds, F128 and F129 because they are the hh motif and contribute a large amount of solvation energy to the interface according to PISA (18), and E140 because it is part of the salt bridges in the interface. We ran co-purification pull-down assays with mutated PPE4_mt_ and EspG_3mt_ (Table 4). As described earlier, EspG_3mt_ is only co-purified with the PE5-PPE4 heterodimer if it forms a complex. The introduction of charges into the buried hydrogen bonds with N127D and N132E was unable to break the PPE4_mt_-EspG_3mt_ interaction, and neither was the charge reversal of E140R, as all three mutations co-purify with EspG_3mt_ (Sup. Fig. 5*a*). This suggests that disruption of any of these single positions is not sufficient to abolish PPE4_mt_-EspG_3mt_ interaction. Conversely, the introduction of charged residues into the hh motif with F128R or F129E did disrupt the interface and prevented EspG_3mt_ from being co-purified (Sup. Fig. 5*a*), as it interrupts with the hydrophobic environment deep within the EspG_3mt_ binding pocket.

**Table 4.** Summary of analysis of PE5_mt_-PPE4_mt_-EspG_3mt_ interactions *in vitro.*

| **PPE4 mutations** |  |
| --- | --- |
| N127D | + |
| F128R | - |
| F129E | - |
| N132E | + |
| E140R | + |
| **EspG<sub>3</sub> mutations** |  |
| R87E | + |
| R102E | + |
| R208E | + |
| E212R | - |
| S231Y | - |

The interface of EspG_3mt_ is also well conserved amongst the various EspG_3_s tested in this study (Sup. Fig. 4), and again, we targeted strictly conserved residues. We selected R208 and E212 because they contain buried hydrogen bonds, R87 and R102 because they form the salt bridge within the interface, and S231 because it sits at the top of the groove of EspG_3_ and could sterically block entrance into the pocket. Neither single mutation of the salt bridge, R87E or R102E, was able to prevent co-purification of EspG_3mt_ (Sup. Fig. 5*b*). Also, the introduction of a charged residue with R208E was unable to prevent the interaction (Sup. Fig. 5*b*). In contrast, E212R was sufficient to prevent co-purification, as well as S231Y (Sup. Fig. 5*b*), as both prevent the hydrophobic tip of PPE4_mt_ from interacting with the binding pocket of EspG_3mt_ either by charge repulsion or steric hindrance. Thus, our mutations on both PPE4_mt_ and EspG_3mt_ highlight the importance of the hydrophobic environment deep within the PPE4_mt_-EspG_3mt_ interface.

### Structure of EspG_3_ in and out of trimer complex

Our structure is the first of EspG_3_ solved in complex with a cognate PE-PPE dimer, and thus we wanted to compare it to the previously solved unbound EspG_3_ structures. In total, there are six available EspG_3_ structures, four of EspG_3ms_ (PDB codes: 4L4W, 4RCL, 5SXL, and 4W4J (13,16)), one EspG_3mt_ (4W4I (13)), and one EspG_3mm_ (5DLB (16)). These six structures can be classified into two different forms, an ‘open’ and a ‘closed’ form. The differentiation between these two forms is the orientation of the C-terminal helical bundle relative to the core β-sheet. The EspG_3mm_ structure (5DLB) is representative of the ‘open’ form, and one of the EspG_3ms_ structures (4RCL) is representative of the ‘closed’ form. Analysis of EspG_3mm_ as it exists in the PE5_mt_-PPE4_mt_-EspG_3mm_ trimer was done relative to these two representative structures. The overall alignment of the representative structures to the bound EspG_3mm_ was good with r.m.s.d. of 2.1 Å and 1.9 Å for the ‘open’ and ‘closed’ forms, respectively (Fig. 4*a*). Inspection of these alignments show the majority of differences to be within the arrangement of the C-terminal helical bundles, with the bound form of EspG_3mm_ being in close to the orientation found in the ‘closed’ form (Fig. 4*b-c*). The bound EspG_3mm_ cannot be any closer to the ‘closed’ form orientation because the C-terminal helical bundle is making contacts with PPE4_mt_. We hypothesized that this C-terminal helical bundle is dynamic and closes on cognate PPE proteins upon interaction. A comparison between the bound EspG_3mm_ structure and the ‘open’ EspG_3mm_ was performed with the DynDom server to test this hypothesis (19). DynDom identified a moving domain within the structures that was located in the C-terminal helical bundle (Fig. 4*d*). DynDom’s analysis also performed a whole structure alignment that agreed with the previous Dali alignment in Fig 4*a-b*. DynDom performed alignments between the fixed domains (residues 11-168 and 189-279) and the moving domains (residues 168-188), which resulted in much better alignments with r.m.s.d. of 1.76 Å and 0.86 Å, respectively. Therefore, the moving domain, the C-terminal helical bundle, is essentially structurally identical between PPE4_mt_-bound EspG_3mm_ and the ‘open’ EspG_3mm_ and its rotation of 30.2° and translation of 0.8 Å is moderately perturbing the fixed domain. Since the moving domain is making extensive contact with PPE4_mt_ and PPE4_mt_ would sterically clash with the current orientation of the C-terminal helical bundle, the movement from the ‘closed’ to the ‘open’ orientation could be significant in releasing the secreted PE-PPE dimers from the chaperone at the secretion machinery.

**Figure 4.**
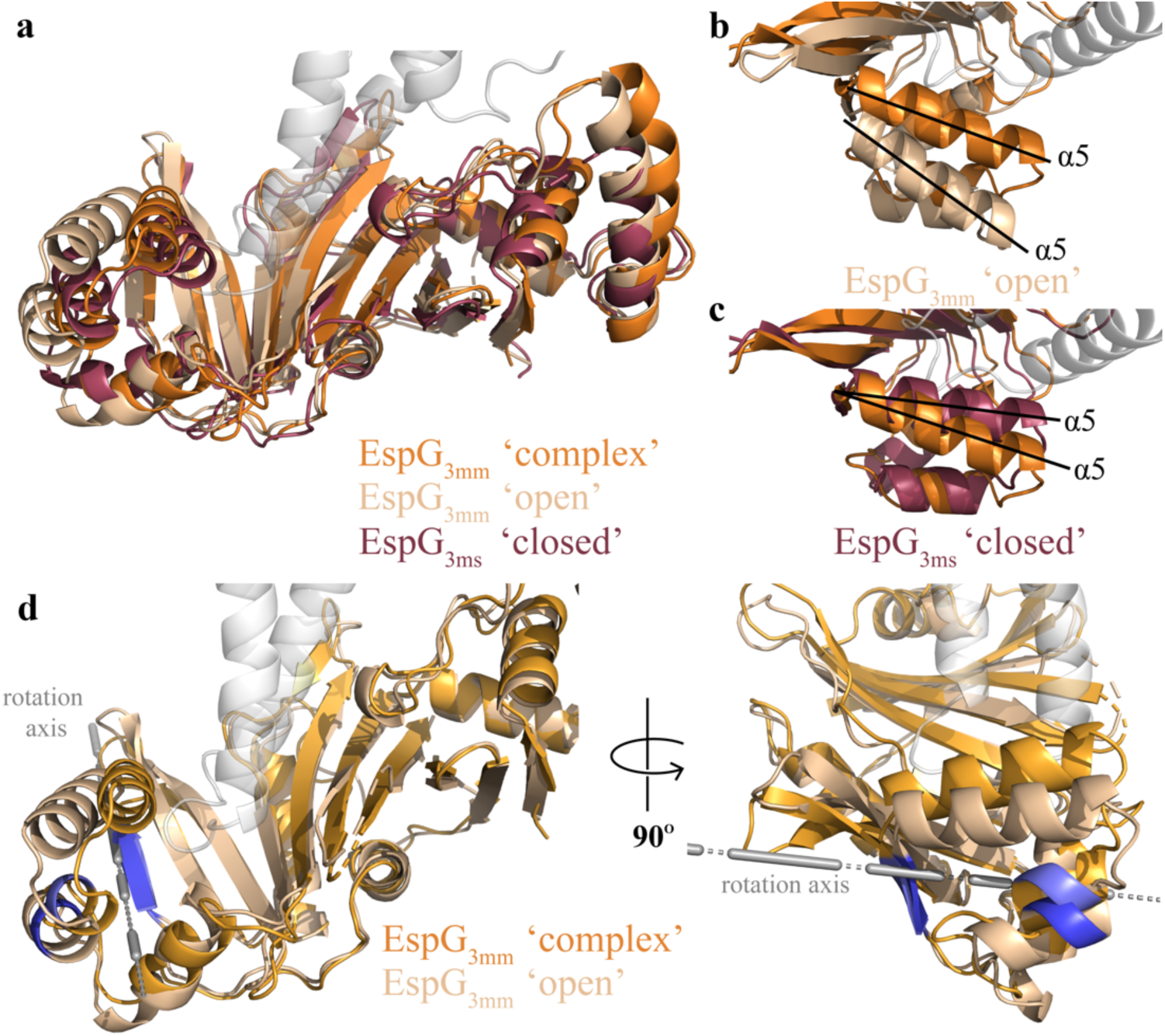
EspG_3_ exists in multiple structural forms. *a*, An ‘open’ (5DLB, EspG_3mm_, sand) and ‘closed’ (4RCL, EspG_3ms_, maroon) conformation of EspG_3_ were aligned to EspG_3mm_ (6UUJ, orange) as it is bound to PPE4_mt_. Overall the different conformations align well to the bound conformation of EspG_3mm_ with r.m.s.d. of 2.1 Å (‘open’) and 1.9 Å (‘closed’). *b and c*, Closeups highlighting the different orientations of the α5 helices in the ‘open’ (*b*) and ‘closed’ (*c*) EspG_3_ structures as compared to EspG_3mm_ bound to PPE4_mt_. *d*, Movement regions defined in EspG_3_ as it moves from the ‘open’ conformation to the bound conformation in two different views related by a 90° rotation. The rotation axis for the moving domain is shown in gray. Each conformation maintains the same coloring as in panel *a*, with the hinge between the moving and fixed domains colored blue (‘open’) and light blue (bound).

### Comparison of ESX-3 and ESX-5 PE-PPE-EspG trimers

A vastly different binding mode is observed when comparing the ESX-3-specific PE5_mt_-PPE4_mt_-EspG_3mm_ trimer to the previously published ESX-5-specific trimers. As mentioned earlier, there is good agreement when comparing individual components of the ESX-3-specific trimer to the available ESX-5-specific trimers (Table 3). The difference between the two sets of trimers became apparent when they were aligned via EspG (Fig. 5*a-b*). Our results focused on comparisons with the PE25_mt_-PPE41_mt_-EspG_5mt_ (4KXR) trimer, but the same differences were present with the PE8_mt_-PPE15_mt_-EspG_5mt_ (5XFS) trimer. The interaction angle of the different PE-PPE heterodimer with EspG is drastically different between the two trimers, with 30° angle difference (Fig. 5*b*). Another difference lies within the hh motif loops of PPE25_mt_ (α4-α5 loop) and PPE4_mt_ (α5-α6 loop) (Fig. 5*c*). In PPE25_mt,_ this loop is seven residues long and undertakes a compact conformation that is not altered during EspG_5mt_ binding (12). In contrast, in PPE4_mt,_ this loop is nine residues long and has an extended conformation, and this difference was rapidly apparent when PPE25_mt_ and PPE4_mt_ were aligned (Fig. 5*c*).

**Figure 5.**
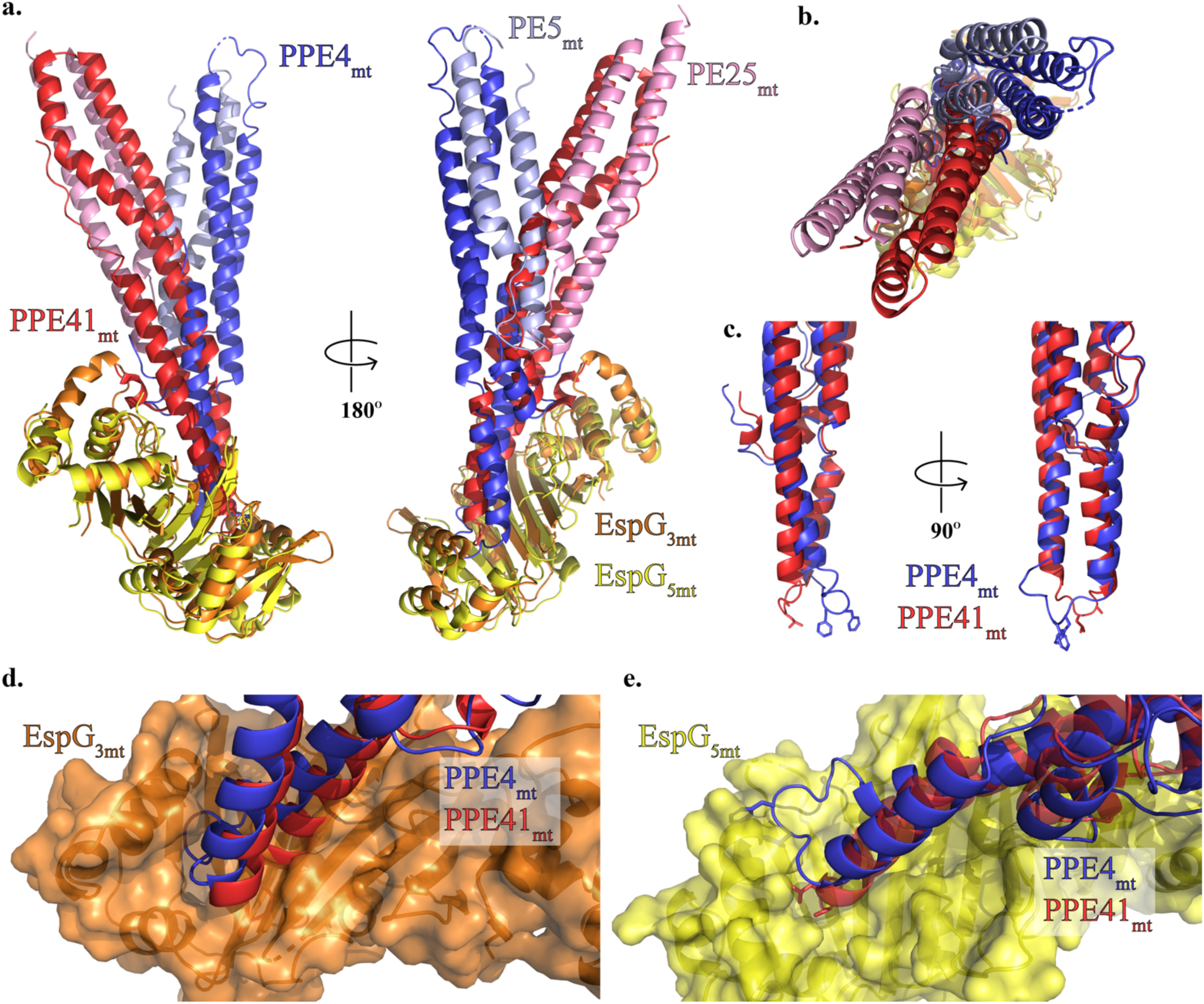
PE5_mt_-PPE4_mt_ interacts with EspG_3mm_ chaperone in a unique mode compared to ESX-5 PE-PPE dimers. *a*, Structural alignment of the ESX-3 and ESX-5 trimers via the EspG chaperones (Dali ref) reveals a difference in the angle of interaction between the PE-PPE heterodimers with their respective chaperone. *b*, Top view of alignment from panel *a. c*, Alignment of PPE41_mt_ and PPE4_mt_ highlights difference in hh loop conformations between ESX-3 (PPE4_mt_) and ESX-5 (PPE41_mt_). *d-e*, Superposition of PPE alignment from panel *c* in context of EspG_3mm_ interaction (*d*) and EspG_5mt_ interaction (*e*) shows the incompatibility of each PPE protein with non-cognate chaperone binding.

This loop conformation also made each PPE protein incompatible with the other’s binding mode. When looking at the PPE alignment in the context of the ESX-3 trimer, the α4-α5 loop of PPE25_mt_ does not align over the central groove of EspG_3mm_ and instead sterically clashes the central β sheet of the chaperone (Fig. 5*d*). The tip of PPE41_mt_ would have to undergo a drastically new tip confirmation in order to bind in the opening of EspG_3mm_. In the context of the ESX-5 trimer, the α5-α6 loop does not align with the central groove of the chaperone, and instead, PPE4_mt_’s hh motif sterically clashes with the C-terminal helical bundle of EspG_5mt_ (Fig. 5*e*). Also, none of the salt bridges between PPE41_mt_ and EspG_5mt_ are conserved in PPE4_mt_. Specifically, D134^PPE41mt^-K235^EspG5mt^, D140PPE41mt-R109EspG5mt, and D144PPE41mt-R27^EspG5mt^; that are all replaced with hydrophobic residues in PPE4_mt_: either T137^PPE4mt^ or L138^PPE4mt^, V144^PPE4mt^, and L147^PPE4mt^, respectively.

## Discussion

In this work, we present the first structure of the PE5_mt_-PPE4_mt_-EspG_3mm_ trimer, which is from the ESX-3 system. Our structure is a mixed trimer, and we presented evidence that EspG_3_ from numerous mycobacterial species can bind the PE5_mt_-PPE4_mt_ heterodimer. Conservation of the EspG_3_s used in this study ranged from 57-83% identity, yet an enrichment in conservation is observed within PPE4-interacting residues (Sup. Fig. 4). The ability of EspG_3_ from numerous mycobacterial species to bind PE5_mt_-PPE4_mt_ suggests that the recognition mechanism is conserved within ESX systems across species. Overall the PE5_mt_-PPE4_mt_ interaction is similar to the previously reported PE-PPE-EspG trimers (12-14) in that PPE4_mt_’s tip is solely interacting with EspG_3mm_ and the general secretion motif of YxxxD/E, on PE5_mt_, is at the distal end of the PE5_mt_-PPE4_mt_ heterodimer. In all copies of PE5_mt,_ this motif is unstructured as it is in the PE8-PPE15-EspG_5_ trimer (14), and similarly, W63^PPE4mt^ is pointed away from this secretion motif. This arrangement is distinct from the PE25-PPE41-EspG_5_ trimers (12,13) and EspB, an ESX-1 substrate that has a similar structural fold to the PE-PPE heterodimers (20,21). PE8_mt_ contains an expanded C-terminal domain, and since the secretion motif is located in the linker between the C-terminal domain and the PE domain the orientation of the secretion motif was unclear (14). PE5_mt_ does not have an expanded C-terminal domain and is just the conserved PE domain, yet its secretion motif is still unstructured in our trimer. Therefore, it is still unclear the exact significance of the structural variations in the ESX secretion motif, and further work is still needed.

Our structure is the first of EspG_3_ bound to a cognate PE-PPE heterodimer. In comparisons of the various published EspG_3_ structures, we identified two different forms that relate to the orientation of the C-terminal helical bundle, an ‘open’ and a ‘closed’ form. EspG_3mm_, when bound to the PE5_mt_-PPE4_mt_ heterodimer, is in a conformation slightly different than the ‘closed’ form due to interactions with the tip of PPE4_mt_. We also found that the C-terminal helical bundle is a dynamic domain and shifts between the ‘open’ and ‘closed’ forms via a hinge movement (Fig. 4*d*). The functional significance of this domain movement could be two-fold. First, the plasticity of the C-terminal helical bundle could allow EspG_3_ to accommodate any variation in the ESX-3-specific PPE tips. While the tip of ESX-3-specific PPE proteins is mostly conserved (Sup. Fig. 3), there is some variations at the end of α5 that could alter the tertiary structure and thus slightly alter the interactions with the EspG_3_ chaperone and the PPE protein. Secondly, the movement of the C-terminal helical bundle could be critical to the release of the PE-PPE heterodimers at the ESX-3 secretion machinery. It is unlikely that PPE4_mt_ could be removed from its interactions with EspG_3mm_ without either movement of the C-terminal helical bundle or steric clashes with the C-terminal helical bundle. Movement of this helical bundle and release of PPE4_mt_ would likely require energy input, and a candidate to provide that energy is EccA. EccA is an ATPase (2) and interacts with both EspG and PPE proteins in yeast two-hybrid experiments (13,22). Recent structures of the ESX machinery from both ESX-3 (4) and ESX-5 (6) suggests overall six-fold symmetry of the core ESX machinery within the inner membrane, and EccA could not only be acting to provide the energy required to uncouple the PE-PPE heterodimers from their EspG chaperone but also to provide a platform for interaction with the core secretion machinery as EccA is likely hexameric when functional.

Previous studies showed that each EspG only recognize PE-PPE heterodimers from their cognate systems (11,12). Despite the structures of two different PE-PPE-EspG trimers from ESX-5 (12-14), it was still unclear how EspG_5_ was differentiating from cognate and non-cognate PE-PPE heterodimers. Our structure represents the first PE-PPE-EspG trimer from ESX-3 and allows for direct comparisons between the ESX-3 and ESX-5 trimers. Our structure reveals that PE5_mt_-PPE4_mt_ interacts with EspG_3mm_ at a different angle of interaction than what was shown for either ESX-5 trimer. This difference in interaction angle presents a different face of PPE4_mt_ to EspG_3mm_. We hypothesize that this is a conserved feature of the ESX-3 PPE-EspG_3_ interaction, as both characterized ESX-5 PE-PPE heterodimers (12-14) display the same face to EspG_5_ despite 33% sequence identity between PPPE41 and PPE15. Therefore, we hypothesize that each ESX system has a unique shape complementarity between its subset of PPE proteins and their cognate EspG chaperone, and these unique shapes are likely not compatiable for interaction with non-cognate chaperones. Our structure is also the first of an ESX-3-specific PE-PPE heterodimer. PE5_mt_-PPE4_mt_ shares the same global conformation as the previously solved PE-PPE heterodimers, yet it differs drastically in PPE4_mt_ in the loop between α5-α6, which contains the hh motif. This longer, more extended loop interacts deeper in the cleft of EspG_3mm_ and is subsequently much more shielded from solvent. It is possible that the longer, extended loop conformation is a feature of ESX-3 PPE proteins and could play an essential role in EspG_3_ recognition.

In conclusion, we presented the first structure of a PE-PPE-EspG trimer from the ESX-3 system. This structure allowed us to compare the interactions of EspG_3_ and a cognate PPE protein to the previously described EspG_5_-PPE interactions. We hypothesize that shape complementarity is a key feature of distinguishing cognate and non-cognate PPE proteins from the EspG chaperones.

## Experimental Procedures

### Bacterial strains and growth conditions

The *Escherichia coli* Rosetta2(DE3) strains grown in Luria-Bertani (LB) medium or on LB agar at 37 °C. When needed, antibiotics were included at the following concentrations: chloramphenicol at 10 μg/ml, streptomycin at 50 μg/ml, and kanamycin at 50 μg/ml.

### Expression and purification of PE5-PPE4-EspG_3_ heterotrimers

Optimized DNA sequences based on the amino acids of full-length PE5 and PPE4 residues 1-180 from *M. tuberculosis* were obtained from Invitrogen and put into a pRSF-NT vector (23) using NcoI and HindIII restriction sites, which contains an N-terminal His_6_ tag on PE5 that is cleavable by TEV protease. EspG_3mt_. EspG_3mm_ expression plasmid was constructed as described previously (12). Mutations in PPE4_mt_, EspG_3mt_, and EspG_3mm_ were introduced with Gibson assembly mutagenesis (SGI-DNA).

Co-expression of all heterotrimers was performed as described previously (12). Briefly, *E. coli* strains containing the appropriate PE5_mt_-PPE4_mt_ and EspG_3_ plasmids were induced with 0.5 mM IPTG when they reached an OD at 600 nm of 0.5-0.8 and then continued to shake at 16 °C for 20 h. Cells were harvested by centrifugation. Cells were then resuspended in lysis buffer (300 mM NaCl, 20 mM Tris pH 8.0, and 10 mM imidazole) and 1:100 Halt protease inhibitor cocktail (Thermo Fisher Scientific, Waltham, MA). Cells were lysed using an EmulsiFlex-C5 homogenizer (Avestin, Ottawa, ON, Canada). The soluble lysate was purified over a Ni-NTA column (G-Biosciences, St. Louis, MO). Eluted protein was dialyzed against lysis buffer without imidazole and incubated with 1:20 mg of TEV protease at 4 °C for 20 h before being re-applied to the Ni-NTA column. Flow-through and washes were pooled and concentrated for size-exclusion chromatography over a Superdex 200 Increase 10/300 GL column (GE Healthcare Life Sciences, Marlborough, MA) that was equilibrated in buffer A (100 mM NaCl and 20 mM HEPES pH 7.5).

### Crystallization, data collection, and structure solution

Purified protein was concentrated to 4.195 mg/ml. Initial screening was done using the MCSG Crystallization Suite (Anatrace, Maumee, OH). This initial screening produced the P2_1_2_1_2_1_ crystals that were grown in 200 mM NH_4_ tartrate and 20% PEG 3350. Optimization around three others hits from the initial crystal screening containing NaCl as the precipitant and various buffers ranging from a pH 5.5 to 8.0 produced the I422 crystals, which were grown in 2.0 M NaCl and 100 mM bis-tris, pH 6.5. Crystals were transferred to cryoprotectant solution, which contained the crystallization solution supplemented with either 20% (P2_1_2_1_2_1_) or 25% (I422) glycerol and then flash-cooled in liquid N_2_. Data were collected at the Southeast Regional Collaborative Access Team (SER-CAT) 22-ID beamline at the Advanced Photon Source, Argonne National Laboratory. Data were processed using *XDS* and *XSCALE* (24). Molecular replacement using Phaser (25) was used to solve the structure of both crystal forms. First, the PE25_mt_-PPE41_mt_ dimer (PDB: 4KXR (12)) and EspG_3mm_ (PDB: 5DLB (16)) were used as search models for the I422 dataset. Later, an early model of the I422 structure was used as a search model for the P2_1_2_1_2_1_ dataset. DENSITY MODIFICATION AND ORIGINAL MODEL. The starting model for both forms was then iteratively rebuilt and refined using Coot and phenix.refine (26,27). The final structure for both crystal forms was refined in phenix.refine, with the P2_1_2_1_2_1_ form using non-crystallographic symmetry restraints (TLS?). All data collection and refinement statistics are listed in Table 1. The final model was assessed using Coot and the MolProbity server (28) for quality.

### Size-exclusion chromatography multi-angle light scattering (SEC-MALS)

Proteins were expressed and purified as described and then passed over an AKTA pure with an inline Superdex 200 Increase 10/300 GL column (GE Healthcare Life Sciences), miniDAWN TREOS, and Optilab T-rEX (Wyatt Technologies, Santa Barbara, CA). The system was equilibrated and run in buffer A. Samples were loaded at a volume of 500 μL at a concentration of 2-4 mg/ml and the system was run at 0.5 mL/min. Analysis of light scattering data was done using Astra (Wyatt Technologies). Molecular weight determination was done by analyzing peaks at one-half their maximum. Graphics were prepared using Prism (Graphpad Software, La Jolla, CA).

### Sequence analysis

Sequence analysis was performed using the EMBL-EBL analysis tools, specifically the Clustal Omega program (29). Rendering of sequence analysis was done with the ESPript server (30).

### Structural analysis

All structural figures were generated using PyMol (http://www.pymol.org) and Chimera/ChimeraX (chimera ref). Electrostatic surface potentials were calculated using the APBS Electrostatics plugin in PyMol (31). Structural alignments were performed with the Dali server (32).

### SAXS Data Comparison and Ab Initio Model Reconstruction

PE5_ms_-PPE4_ms_-EspG_3ms_ trimer SAXS data (SASDDX2 (16)) was compared to a single copy of the mixed PE5_mt_-PPE4_mt_-EspG_3mm_ trimer structure (PDB code: 6UUJ) using CRYSOL (17). Ab initio reconstruction of the envelope was completed using GASBOR (33). Monomeric symmetry was used as a constraint for GASBOR. Twenty ab initio models were generated and averaged using the DAMAVER software package (34). DAMSEL rejected only one model.

### Accession codes

Coordinates and structure factors were deposited in the Protein Data Bank with accession codes 6UUJ (P2_1_2_1_2_1_) and 6VHR (I422).

## Acknowledgments

The work in this study was supported by an Institutional Development Award (IDeA) from the National Institute of General Medical Sciences of the National Institutes of Health grants P20GM103486 and P30GM110787, and by the National Institute of Allergy and Infectious Diseases grant R01AI119022 to K.V.K. W.A.C. was supported by the National Science Foundation Research Experiences for Undergraduates (REU) grant 1358627. Use of SER-CAT is supported by its member institutions (see www.ser-cat.org/members.html), and equipment grants (S10_RR25528 and S10_RR028976) from the National Institutes of Health. Use of the Advanced Photon Source was supported by the U. S. Department of Energy, Office of Science, Office of Basic Energy Sciences, contract W-31-109-Eng-38. The contents of this publication are solely the responsibility of the authors and do not necessarily represent the official views of NIGMS or NIH.

