## Supporting Information for "PE5-PPE4-EspG_3_ trimer structure from mycobacterial ESX-3 secretion system gives insight into cognate substrate recognition by ESX systems"

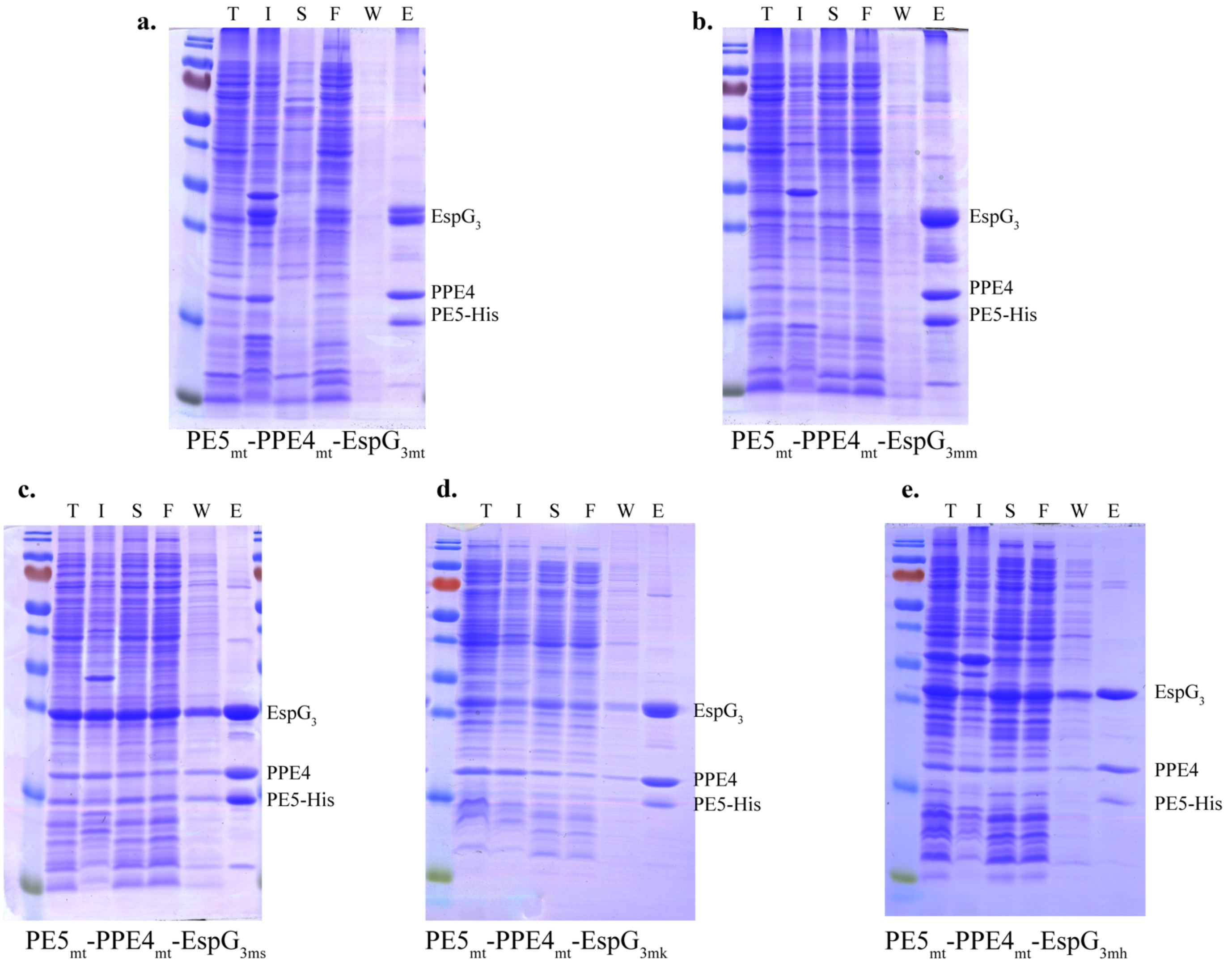


**Supplementary Figure 1. PE5$_{mt}$-PPE4$_{mt}$ dimer is bound by EspG$_3$ from various mycobacterial species.** Co-purification of PE5$_{mt}$-PPE4$_{mtb}$with *a,* EspG$_{3mt}$, *b,* EspG$_{3mm}$, *c,* EspG$_{3ms}$, *d,* EspG$_{3mk}$, or *e,* EspG$_{3mh}$. T is total lysate, I is insoluble lysate, S is soluble lysate, F is column flow through, W is column wash, and E is column elution.

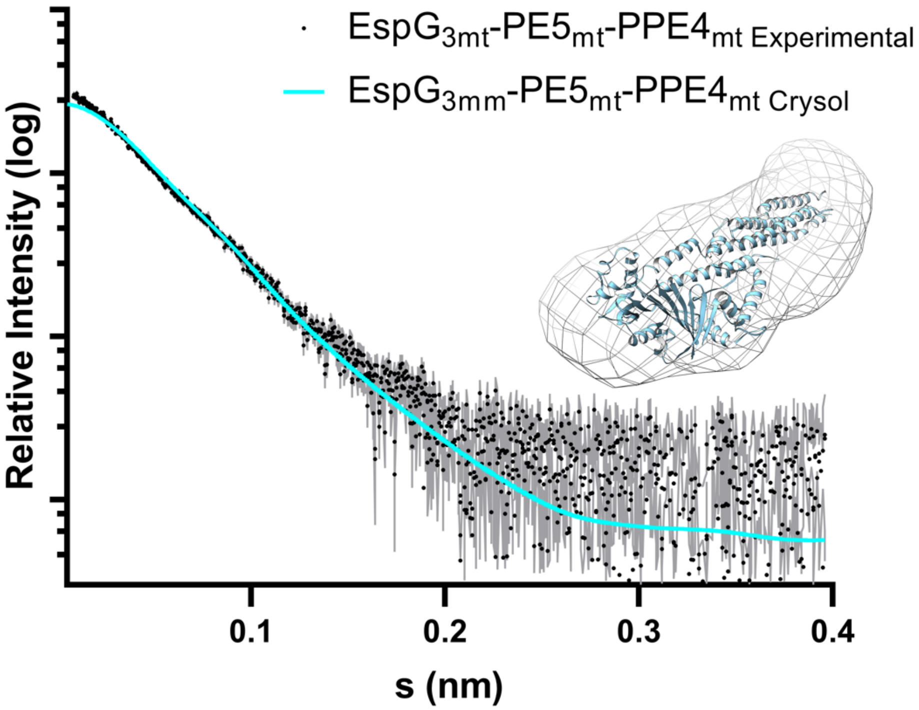


**Supplementary Figure 2. Comparison of $PE5_{mt}$-$PPE4_{mt}$-$EspG_{3mm}$ crystal structure and $PE5_{ms}$-$PPE4_{ms}$-$EspG_{3ms}$ SAXS data.** The SAXS data was originally collected in (ref) and compared to the 6UUJ structure we obtained. The $\chi^2$ between the crystal structure and SAXS data is 2.53 as compared by CRYSOL (ref). An insert shows the 6UUJ structure inside an envelope created by GABSOR (ref).

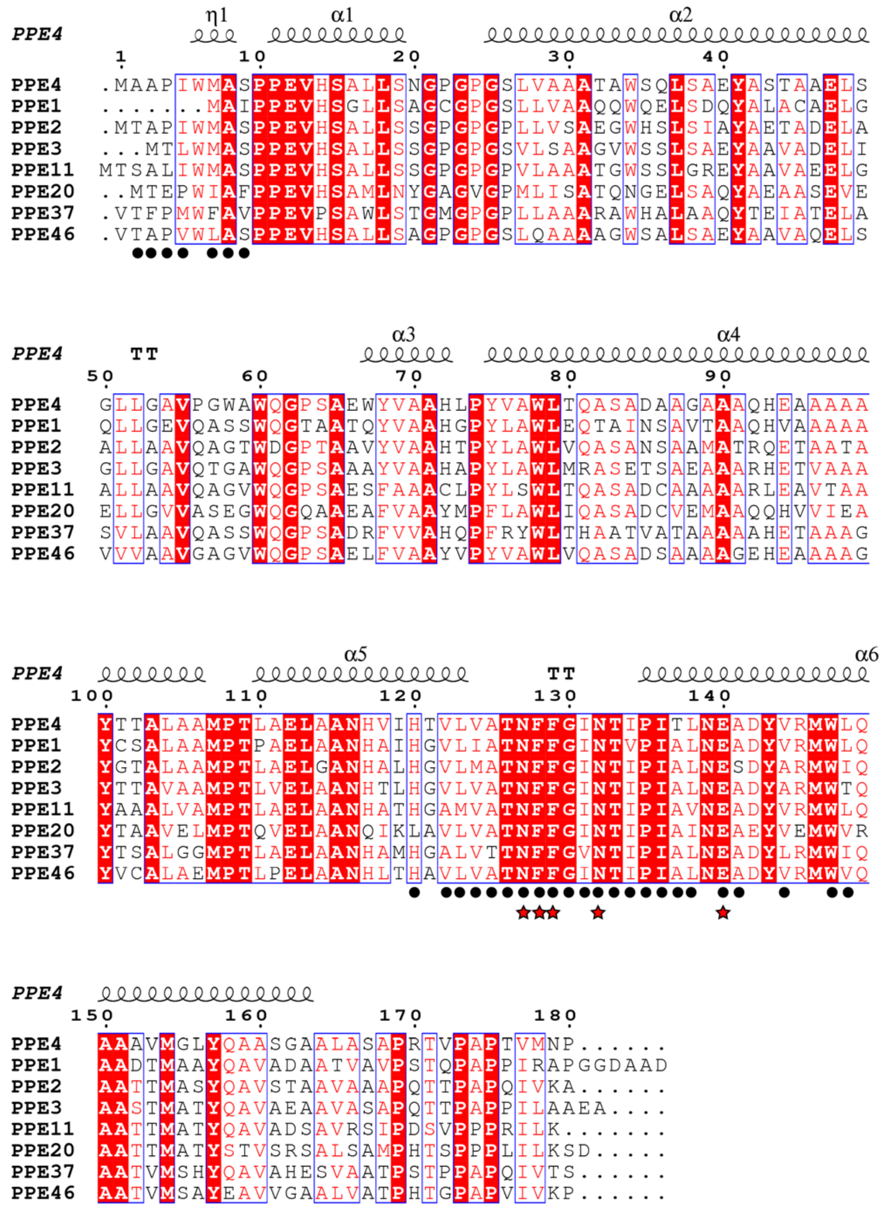


**Supplementary Figure 3 (FINAL). Sequence alignment of *M. tuberculosis* ESX-3-specific PPE genes.** Genomic sequences of *M. tuberculosis* PPE proteins were aligned with Clustal (ref). The secondary structure from $PPE4_{mt}$ from our trimer model (6UUJ) is shown above each row of the alignment. Residues that are identical across all of the ESX-3-specific PPEs are highlighted in red. Residues interacting with $EspG_{3mm}$ in the crystal structure are denoted with black circles and the ones that were chosen for mutagenesis are denoted with a red star.

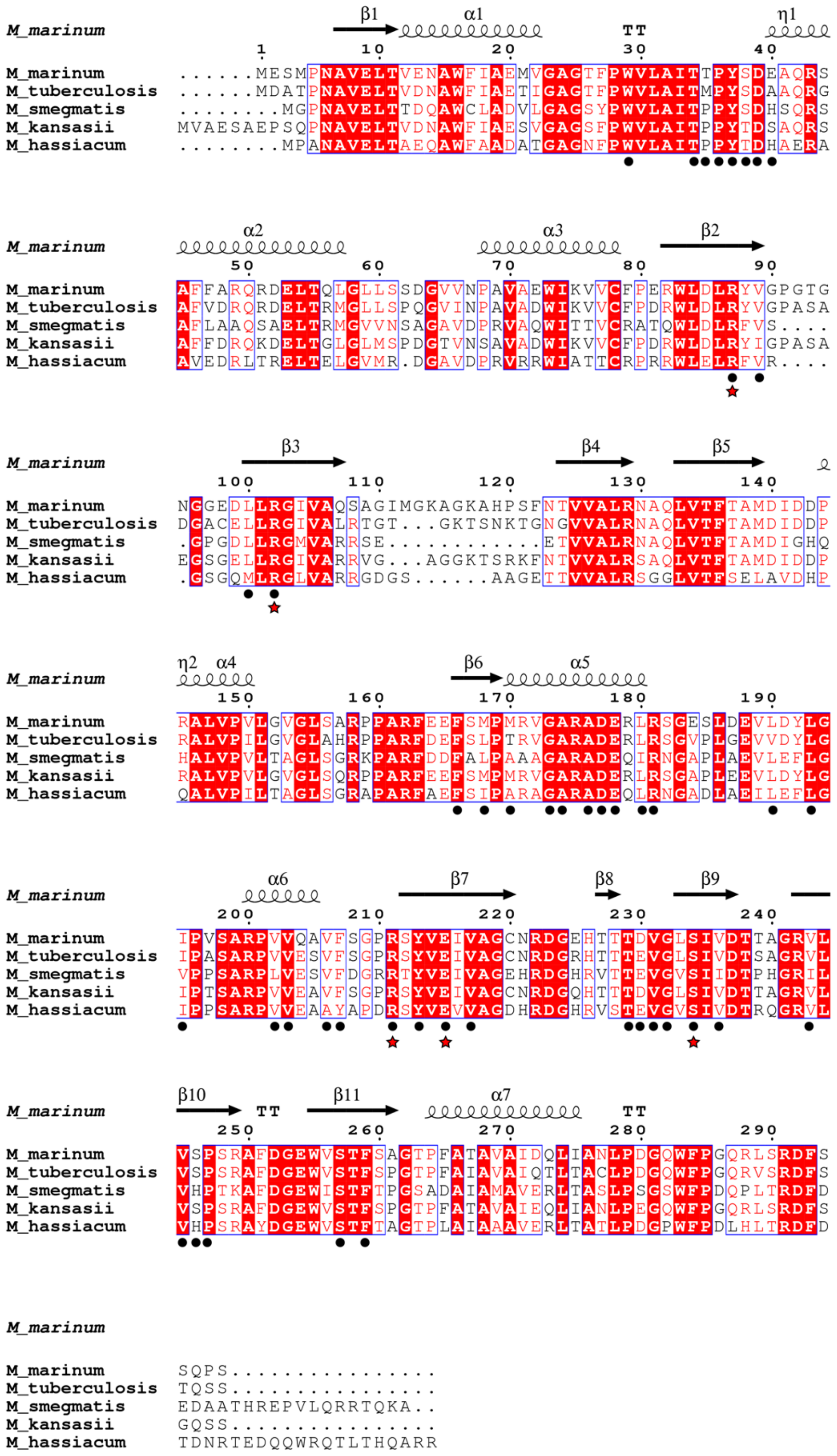


**Supplementary Figure 4 (FINAL). Sequence alignment of selected EspG$_3$'s shows interacting residues are conserved.** Genomic sequences of the five EspG$_3$s used in this study were aligned using Clustal (ref). The secondary structure from EspG$_{3mm}$ from our trimer model (6UUJ) is shown above each row of the alignment. Residues that are identical across the five species are highlighted in red. Residues interacting with PPE4$_{mt}$ in the crystal structure are denoted with black circles and the ones that were chosen for mutagenesis are denoted with a red star.

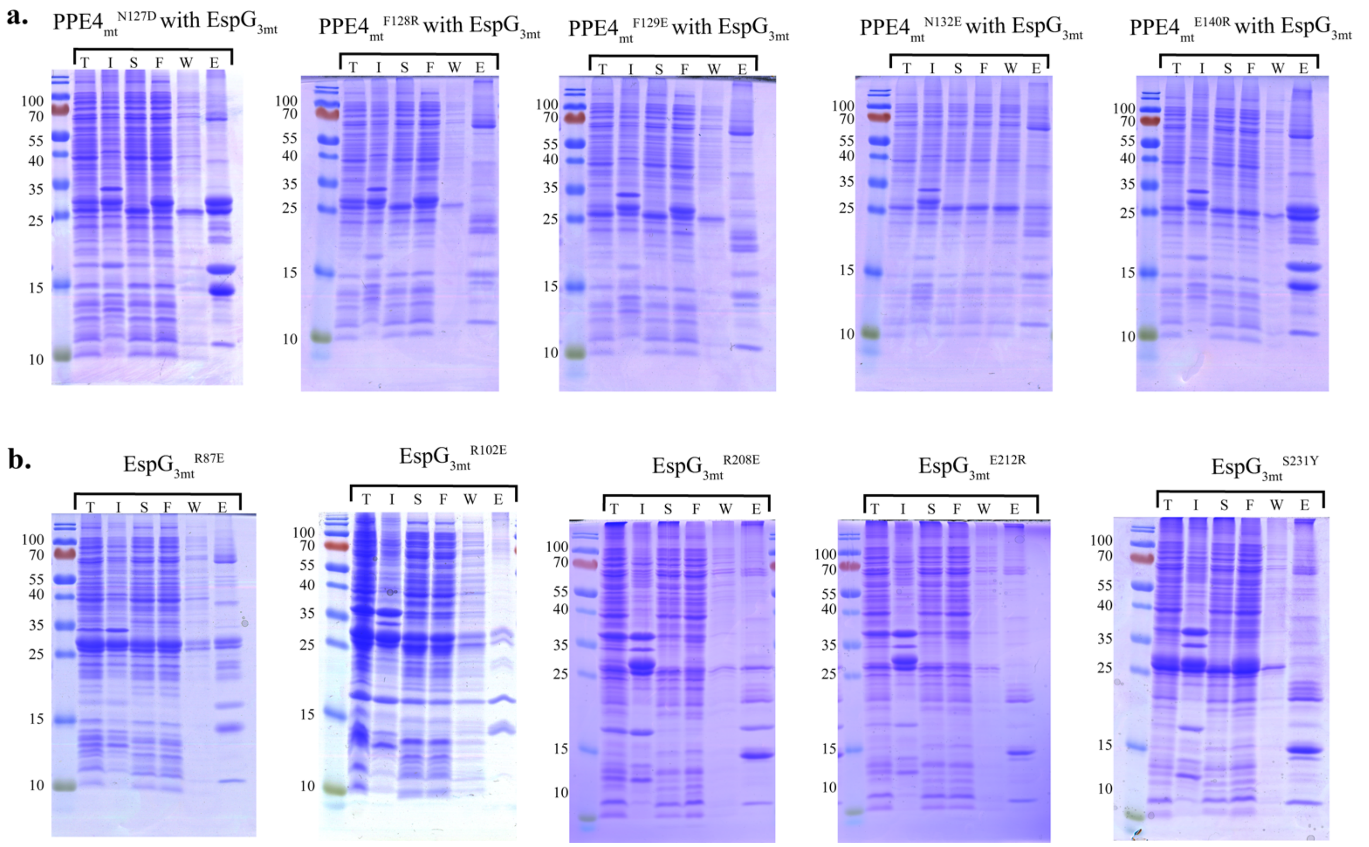


**Supplementary Figure 5 (FINAL). Co-purification of selected PPE4$_{mt}$ and EspG$_{3mt}$ mutants with their wild-type partners.** Gels from co-purification pulldowns of PPE4$_{mt}$ (*a*) and EspG$_{3mt}$ (*b*) mutations show which mutations disrupt the PPE4$_{mt}$-EspG$_{3mt}$ interface (PPE4$_{mt}^{F128R}$, PPE4$_{mt}^{F129E}$, EspG$_{3mt}^{E212R}$, EspG$_{3mt}^{S231Y}$), and which do not. Results are summarized in Table 2. T is total lysate, I is insoluble lysate, S is soluble lysate, F is column flow through, W is column wash, and E is column elution.

**Supplementary Table 1.** Summary of constructs utilized for crystallization experiments and final outcomes.

| **PE5 construct (all constructs contain $His_6$ purification tag)** | **PPE4 construct (only N-terminal PPE domain)** | **$EspG_3$ construct** | **Crystallization Outcome** |
|---|---|---|---|
| *MSMEG_0618* | *MSMEG_0619* | *MSMEG_0622* | Low resolution crystals |
| *MSMEG_0618* with MBP fusion (two different linker lengths) | *MSMEG_0619* | *MSMEG_0622* | Low resolution crystals for both linker lengths |
| *MSMEG_0618* with T4L fusion (3 different forms of T4L) | *MSMEG_0619* | *MSMEG_0622* | Poor expression of trimer in all forms |
| *Rv0285* | *Rv0286* | *Rv0289* | Low resolution crystals |
| *Rv0285* | *Rv0286* | *MMAR_0548* | Two crystal forms solved |
